# Active vision modulates early cortical stages of visual processing: Evidence from MEG and eye-tracking

**DOI:** 10.1101/2025.11.17.688782

**Authors:** Christoph Huber-Huber, Floris P. De Lange

## Abstract

Vision is fundamentally active. Humans and other mammals make several saccadic eye movements per second. This feature of mammalian vision has received increasing interest across the last decades and many behavioural and neurophysiological studies have investigated how the visual system accounts for the effects of self-generated motion (efference copy) in perceptual processing. Still, it is often believed that, besides accounting for the efference copy and for the rhythmic modulation of activity due to perpetual saccade-fixation cycles, active vision does not further modulate neural processing of visual input obtained during periods of fixation. In contrast, using a gaze-contingent trans-saccadic change design with concurrent MEG and eye-tracking data recording from human participants, we find sequential history effects in visual cortices. Harnessing the preview effect, we show that primary visual and ventral occipital cortices respond differently when eye-movements bring objects into focus compared to when the same objects appear during steady fixation. Specifically, we find that sequential modulations of cortical processing across saccades, in the form of preview effects, are increased during active vision compared to steady-gaze control conditions. Our data also show that these sequential modulations cannot be explained by any type of adaptation alone. Our study, therefore, provides an update on the role of active vision in modulating cortical function.

## Introduction

Active vision has been an upcoming research topic across the last decades ^1–3^. Humans make about three to four saccadic eye movements per second in daily life and it is clear that the visual system accounts for the perceptual effects of these self-generated actions – the corollary discharge based on an efference copy ^4,5^ – to arrive at a stable visual percept ^6,7^. While ideas of active vision can be traced back in history ^8,9^, contemporary research provides increasing empirical evidence for the interplay between eye movements on perceptual processing in humans ^10,11^ and for the flexible adjustment of these processes according to sensorimotor contingencies ^12,13^.

Still, it is often believed that, besides accounting for the efference copy, active vision does not any further modulate neural processing of visual input obtained during the fixation periods of perpetual saccade-fixation cycles. The logic here is that any saccade-locked or fixation-locked neural response represents the sum of oculomotor activity and the actual change in visual stimulation elicited by a saccadic eye movement. In order to isolate the oculomotor contribution, that is, the contribution of active vision, one has to compare two conditions with identical visual input that only differ in whether a saccade has been made, which usually means to compare an active saccade condition to a condition with saccade-like stimulation under stable fixation. Investigating the responses of neurons in V1 and further downstream in the ventral visual cortices of non-human primates to stimuli that appear in the visual field because of a saccade and contrasting them to saccade-like simulated stimulus onsets has suggested no significant modulation by active vision ^14–17^.

The findings of similar responses in V1 and ventral visual cortices for real saccades compared to saccade-like stimulation are contrasted by two exceptions ^18,19^, but for these two studies the purported effect of active vision can in theory be explained by differences in visual and intra-saccadic stimulation ^3,10,11,20,21^. MacEvoy and colleagues contrasted a condition in which the monkey made a saccade to a small bar stimulus with a condition in which the same stimulus did not appear before 500 ms after saccade offset, which means that there was an apparent difference in visual stimulation between these two conditions ^18^. McFarland and colleagues directly compared a condition in which monkeys made saccades on natural scene images to a condition in which the images were rapidly translated while the monkey maintained fixation ^19^. However, the image translation achieved with a 100 Hz monitor is not comparable to the analogue visual transient created by a real saccadic eye movement and, thus, the modulation of neural activity in the saccade compared to the control condition is confounded with the effect of visual transients on post-saccadic neurophysiological activity ^10,22,23^.

In addition to these limitations, previous studies on neural responses in visual cortices largely ignored the availability of pre-saccadic preview information which is inherent to active vision ^14,24,25^ and if they did simulated pre-saccadic previews, they did not find much of a difference between active and passive vision conditions. For instance, Vinje and Gallant accounted for the complete dynamics of active vision through a passive saccade-simulation condition including simulated previews, however, they examined particular types of neural coding principles which did not appear to differ much between active and passive vision conditions ^26^. Another study used a free-viewing paradigm which, per definition, includes pre-saccadic previews, but they did not find much evidence that saccade execution affected neural activity in V1 or further downstream in ventral cortices apart from initiating a feedforward processing cascade ^27^. In addition, they missed a passive vision control condition.

Some evidence for a modulation of neural activity in V1 and other visual brain areas by active eye movements comes, however, from research on active sensing and active sampling ^28–30^. A central finding of this line of research is the observation of greater neural excitability after fixation onsets, usually around 50-100 ms, when contrasted to other periods within a saccade-fixation cycle ^31^ or to a pre-fixation baseline ^32^. Importantly, this greater excitability exhibits a laminar profile distinct from the feedforward processing observed during stable fixation. Still, albeit highlighting the rhythmic nature of neural activity timed with eye movements, these modulations have not yet been contrasted to passive-viewing control conditions that mimic the visual input elicited by saccades and that would allow to attribute the modulation with certainty to active vision. In principle, the reported modulations could be attributed to the specific spatiotemporal dynamics of visual input that are generated by eye movements ^10,21^.

Previous research has, thus, not yet provided compelling evidence for the idea that the visual cortical processing during the time of a fixation is influenced by active vision, apart from exhibiting rhythmic modulations timed to eye movements. However, it seems unlikely that visual cortices would not show any further modulation by active vision in terms of sequence effects or effects of viewing history given the clear influence of prior extrafoveal information on post-saccadic foveal processing ^33^. This influence has been demonstrated with preview effect paradigms in which visual input at the future saccade target location is changed in a gaze-contingent way during saccade execution. Such trans-saccadic changes (invalid previews) result in behavioral performance decrements in perceptual discrimination tasks and in larger fixation-related evoked responses in the electroencephalogram (EEG) compared to no-change (valid preview) conditions ^34–37^. Preview effects are also well studied in the context of reading research where they provide insights in how much information readers can extract parafoveally from upcoming words ^38–42^. Although preview effects have been shown to be larger during active saccade compared to passive viewing conditions ^43,44^, which suggests that they are related to active vision, it is still unclear to what extent such trans-saccadic sequential effects are supported by visual cortices, given the low spatial precision of EEG and the lack of compelling evidence from monkey research for sequential history effects specific to active vision in these brain areas. Moreover, recent results on preview effects in the superior colliculus ^45^ rather suggest that these sequential effects are actually supported by subcortical brain structures.

Using source localized magnetoencephalography (MEG) and eye-tracking coregistration, in the present study, we investigated whether visual cortices show sequential viewing history effects across saccades that are indeed specific to saccade execution, that is, specific to active vision. Our experiment consisted of two types of active and one passive viewing condition, administered in separate blocks. In the *saccade* viewing blocks, participants actively made saccades to an extrafoveally presented object which could change (invalid preview) or remain the same (valid preview) during the saccade. In the *replay* viewing blocks, participants kept their gaze fixed at the center of the screen and instead of moving their eyes, the object moved to the center of the screen. This condition mimicked the visual input obtained in the saccade blocks as well as possible without actual saccade execution. Crucially, all viewing blocks contained the same number of valid (no trans-saccadic stimulus change) and invalid (trans-saccadic change) preview trials, which allowed us to attribute any differences in the preview effect contrast (invalid minus valid) between saccade and replay blocks to active vision instead of to any residual visual stimulation discrepancies that might have still existed between saccade and replay blocks, because any such discrepancies are cancelled out in the preview effect contrast. In order to equate the conditions also in their temporal predictability and because the gaze-contingent target onset in the saccade condition is predictable from saccade execution, in the replay condition object motion was played-back after a fixed time delay for each participant. If visual cortices exhibit sequential viewing history effects due to active vision we should find at least a larger preview effect in the saccade compared to the replay blocks in regions of interest (ROI) covering the visual cortex.

A common explanation for a change in neural activity in response to repeated stimulation (valid preview) compared to a change in stimulation (invalid preview) is neural adaptation ^46^. To rule out that neural adaptation confounded the expected modulation of cortical activity by active vision, we included a second type of active viewing blocks, the *blank* blocks, which were the same as the saccade blocks except that we presented a blank screen for 400 ms before the target onset. This manipulation was included because, besides retinotopic adaptation, there is evidence for specifically spatiotopic adaptation and the pre-saccadic preview and post-saccadic target stimuli in our design occur at exactly the same spatiotopic location in the saccade blocks, but not in the replay blocks, making spatiotopic adaptation a likely confound with any modulation by active vision. Crucially, however, spatiotopic neural adaptation is known to increase with a blank interval between adaptor and test stimuli ^47–49^ ^but^ ^see^ ^50^, which means that, if spatiotopic adaptation was responsible for the greater preview effect in the saccade compared to the replay blocks, we should observe an even greater preview effect in the blank blocks than in the saccade blocks.

Our experimental design contained active eye movements or simulated replay equivalents and gaze-contingent trans-saccadic changes which means that one trial contains more than one stimulus onset with potential follow-up saccades and fixations. Each event triggers an MEG-response and to separate these temporally overlapping responses we used temporal deconvolution ^51,52,see^ ^also^ ^53,54^. This method provides the additional advantage of being able to explicitly model any systematic variation in gaze behavior across conditions which we had observed as well.

To preview our results, we found a more pronounced difference between valid and invalid preview conditions in the saccade blocks than in the replay blocks and no preview effects in the blank blocks in primary visual (V1) cortex. This result shows that early visual cortex exhibits more pronounced trans-saccadic viewing history effects when participants make saccades, compared to when the same visual input is provided without making a saccade.

## Results

### Viewing history effects in eye movements and behavior necessitate temporal deconvolution of MEG data

To investigate whether visual cortices show effects of sequential viewing history across saccades (preview effects), which are indeed specific to saccade execution, we measured the source-localized MEG signal time-locked to target foveation in three different types of viewing blocks (Figure 1). However, the preview conditions and viewing blocks did not only affect the MEG signal but also the participants’ eye movements. In the saccade and blank blocks, where participants made a cued saccade to a pre-saccadic object (valid preview) or to a phase-scrambled and blurred version of the same object (invalid preview) (Figure 1A, 1C, 1D), the saccade amplitudes were minimally (by 0.22°), but very consistently, greater for valid (8.13°) than for invalid previews (7.91°) (F(1,35) = 39.71, p < .001, Figure 1F). We, therefore, included each trial’s saccade amplitude as covariate in an MEG temporal deconvolution model to explicitly capture the effect of saccade amplitudes on the first post-saccadic EEG/MEG component, aka the lambda response ^51,52,55^, because the lambda response extends into the time window of 100-300 ms where preview effects are usually found and where differences in saccade amplitudes between conditions might otherwise confound conclusions from the MEG signal.

**Figure 1.**
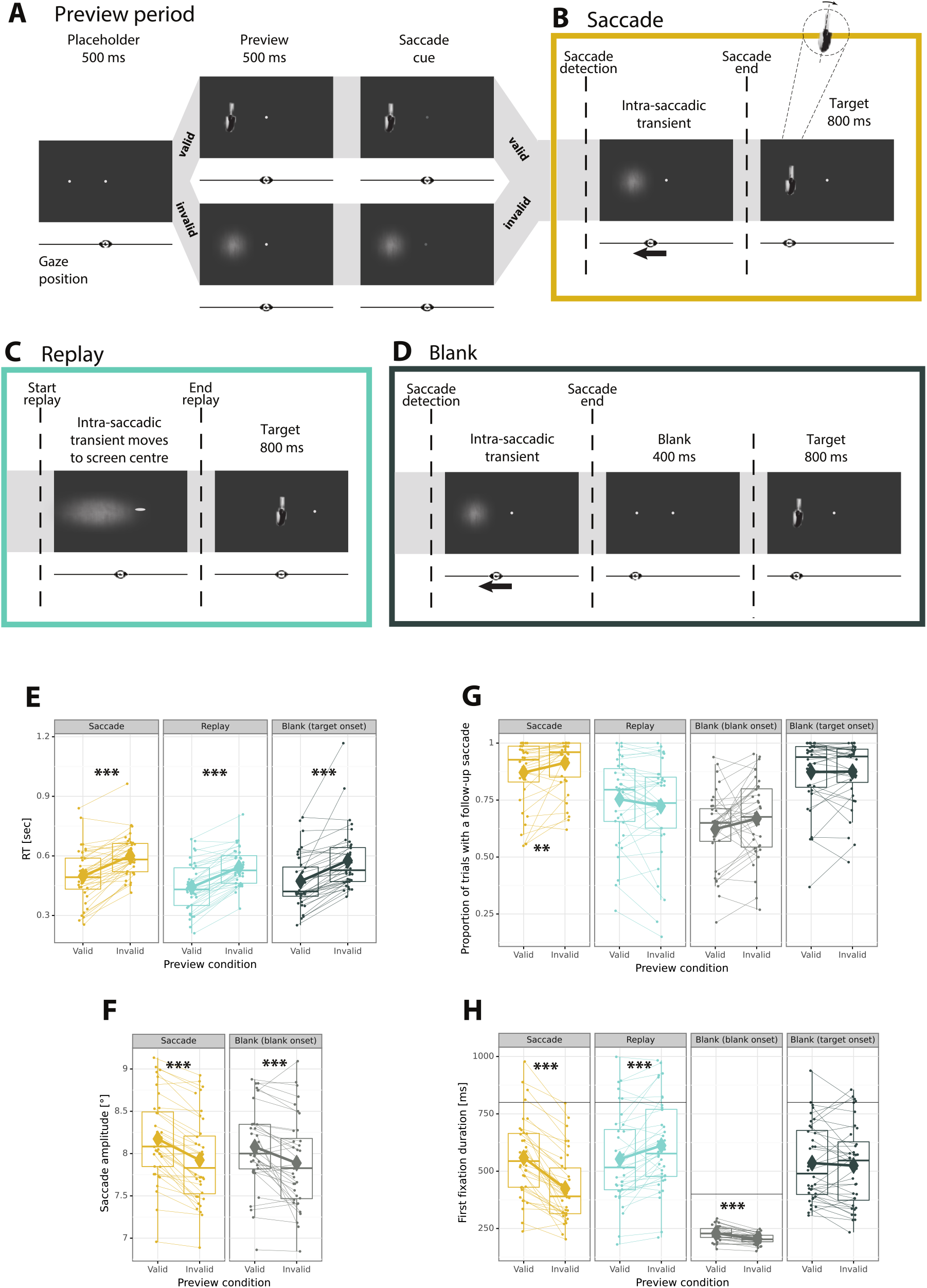
Trial procedure and behavioral results. (A) Stable fixation for 500 ms triggered the presentation of one out of 80 objects, here a garden shovel (valid preview) or their blurred version (invalid preview). In the saccade blocks (B), the participant made a cued saccade to that preview object which appeared after the saccade as intact, non-blurred target object in both the valid and invalid preview conditions. In the replay blocks (C), the participants maintained stable fixation throughout the trial and the object moved into the foveal visual field after a saccadic latency determined per participant depending on their practice trial performance. The blank blocks (D) were the same as the saccade blocks except that a blank screen was presented for 400 ms before the target onset. In all three viewing conditions, participants reported whether the final target object was tilted left or right. (E) Manual response times showed a significant preview effect in all three viewing conditions. (F) Saccades were significantly larger with valid than with invalid preview for both saccade and blank viewing. (G) In the saccade blocks, the proportion of trials with a follow-up eye movement after the initial target fixation was larger with an invalid compared to with a valid preview. (H) In the saccade blocks, the first fixation on the target (or blank) was longer with a valid than with an invalid preview. This effect was the same for fixations on the blank screen, was reversed for the replay blocks, and was gone after the final target onset in the blank blocks. The black horizontal line indicates target stimulus offsets (800 ms) in all blocks and the end of the blank screen (400 ms) for the blank onset.

Interestingly, the preview conditions did not only affect saccade amplitudes but also the follow-up gaze behavior after the initial target fixation and, crucially, this effect varied across viewing blocks. With an invalid preview, participants were more likely to make a follow-up saccade than with a valid preview (Figure 1G), but only in the saccade block (t(35) = 3.11, p < .010). In the replay block, this effect was reversed (interaction F(1,35) = 11.04, p = .002). In the blank block, the effect was comparable to the saccade condition (interaction F(1,35) = 0.01, p = .912) considering first the onset of the blank screen. The subsequent target onset did not appear to further modulate follow-up saccades (interaction with saccade condition, F(1,35) = 6.44, p = .016, invalid 875 ms, valid 873 ms, t(35) = 0.14, p = .890). If a follow-up saccade was made, the time until this saccade happened, that is the duration of the first fixation on the target (Figure 1H), showed the opposite pattern in the saccade (t(35) = 9.56, p < .001) compared to the replay block (t(35) = 4.21, p < .001, interaction F(1,35) = 107.99, p < .001). In the blank blocks, there are two first fixation onsets because of the intermediate blank screen, one on the blank screen and one on the target. For the blank screen onset, the timing of follow-up saccades was similar as in the saccade blocks but less extreme (interaction F(1,35) = 62.50, p < .001). After the onset of the target, which followed the 400-ms blank screen, there was no evidence anymore for an effect of the preview (t(35) = 0.79, p = .436, interaction with saccade blocks F(1,35) = 54.77, p < .001). Because of these patterns in eye movements, we included follow-up saccades as additional events in our deconvolution model. This allowed us to separate the MEG response to the initial fixation from the response elicited by follow-up saccades ^51,52^.

After making a saccade to the extrafoveally presented objects (Figure 1B and 1D) or watching the replay (Figure 1C), participants had to report in each trial whether the object was tilted left or right (Figure 1). As expected, this manual response showed a preview effect (Figure 1E). Response times were faster with a valid (471 ms) compared to an invalid (573 ms) preview (F(1,35) = 116.64, p < .001) and there was no evidence for a modulation by the viewing blocks (both interaction p > .823). Some participants, however, showed on average very early response times, even within the time period of a neural preview effect around 200-250 ms (cf. Figure 1E). In order to avoid that these occasionally very early responses could confound our conclusions from the MEG signal, we added manual responses as separate events to our temporal deconvolution model. Error rates did not provide any evidence for preview effects or differences between viewing blocks (all p > .076).

### Active vision modulates viewing history effects in the early visual cortex

To investigate whether active vision modulates the impact of the pre-saccadic preview on processing the post-saccadic target in visual cortices, we source-localized the MEG signal to five visual cortical surface ROIs per hemisphere: V1, V2, V34, ventral occipital (VO) because of its involvement in spatiotopic adaptation ^48^, and lateral occipital (LO) because of its involvement in object processing and sensory predictions ^56^. To capture potential effects related to the efference copy and saccadic remapping, we added two parietal areas: an intraparietal ROI termed IP ^57,58^, and the supramarginal gyrus (SMG) ^59^ (Figure 2A). At each ROI, we extracted the MEG response time-locked to foveating the target object at the participant-level using temporal deconvolution ^52^ for each preview (valid, invalid) and each viewing block condition (saccade, replay, blank). Except for left SMG and left VO the deconvolution models’ R2 were significantly greater than their cross-validated counterparts, confirming that, despite their complexity, the models were not overfitting the data for any of the ROIs that showed significant experimental effects later on (Figure 2B).

**Figure 2.**
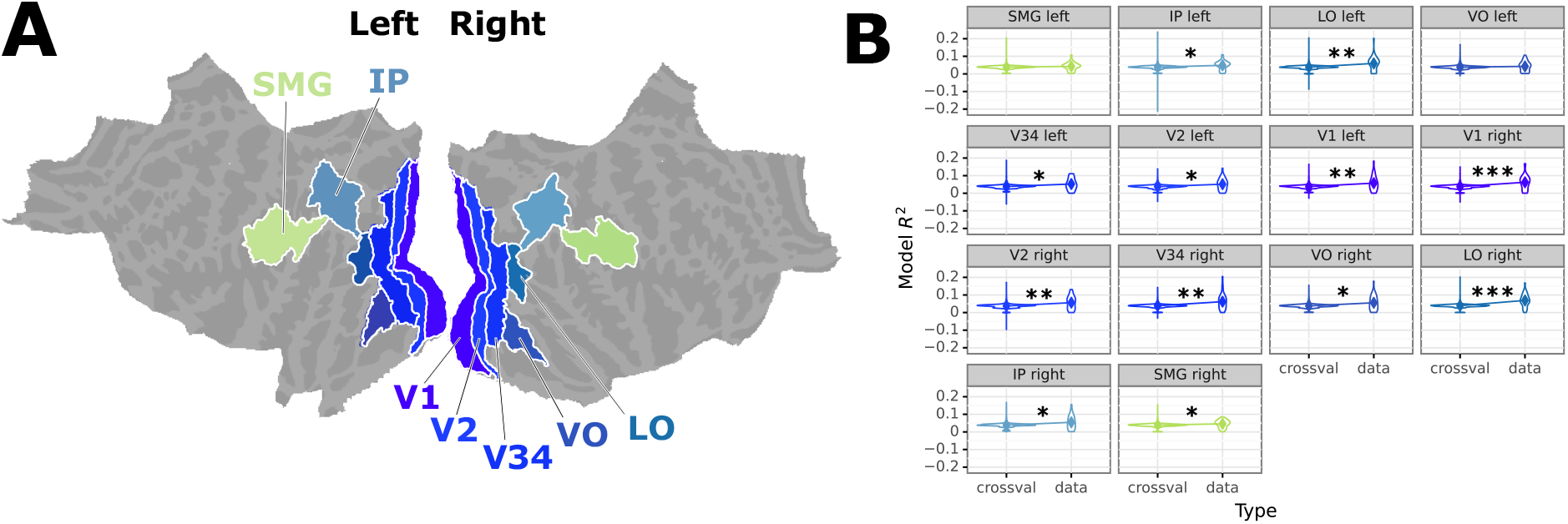
**(A)** MEG source regions of interest (ROI) based on Glasser et al.^60^ illustrated on the fsaverage freesurfer brain. We included all visual areas V1, V2, and V3 plus V4 merged into area V34 in order to exhibit a similar number of vertices compared to V1 and V2. In addition, we defined two mid-higher level visual areas: a lateral occipital area (LO) and a ventral occipital area (VO), and two parietal areas because of their involvement in trans-saccadic processing: an intraparietal area (IP) and the supramarginal gyrus (SMG). ROIs for the left and right hemispheres were kept separate to capture any lateralized visual activity due to the lateralized preview stimulus. **(B)** Deconvolution model fit R2 of the original model (data) compared to 100 crossvalidation runs with scrambled event information (crossval). Asterisks indicate significantly better (within-participants paired t-tests) fit for the original compared to the crossvalidation data for all ROIs except for left SMG and left VO. For further details see section Materials and Methods, MEG and eye-tracking data processing, Regions of interest.

As expected, we found a clear preview effect (invalid – valid) indicating an impact of the viewing history across saccades in right V1 and this effect was gone in the passive replay condition (Figure 3A and 3B, planned-contrast interaction effect clusters for viewing (saccade, replay) x preview (valid, invalid) illustrated by the horizontal bar in Figure 3D). This result shows that active vision leads to an impact of previously obtained input from the saccade target location which is absent in passive viewing. To corroborate the spatial specificity of the preview effect in V1, we examined the source leakage of right V1 ROI to all other single source dipole locations of the whole cortical surface ^61^. As illustrated in Figure 3E, the source leakage from and to V1 was constricted to occipital sources largely within the defined ROI. Moreover, note that the significant cluster for the preview effect in the saccade blocks (Figure 3A) matches the time period of preview effects in the EEG, which is around 200 ms after fixation onset ^34–36,39^.

**Figure 3.**
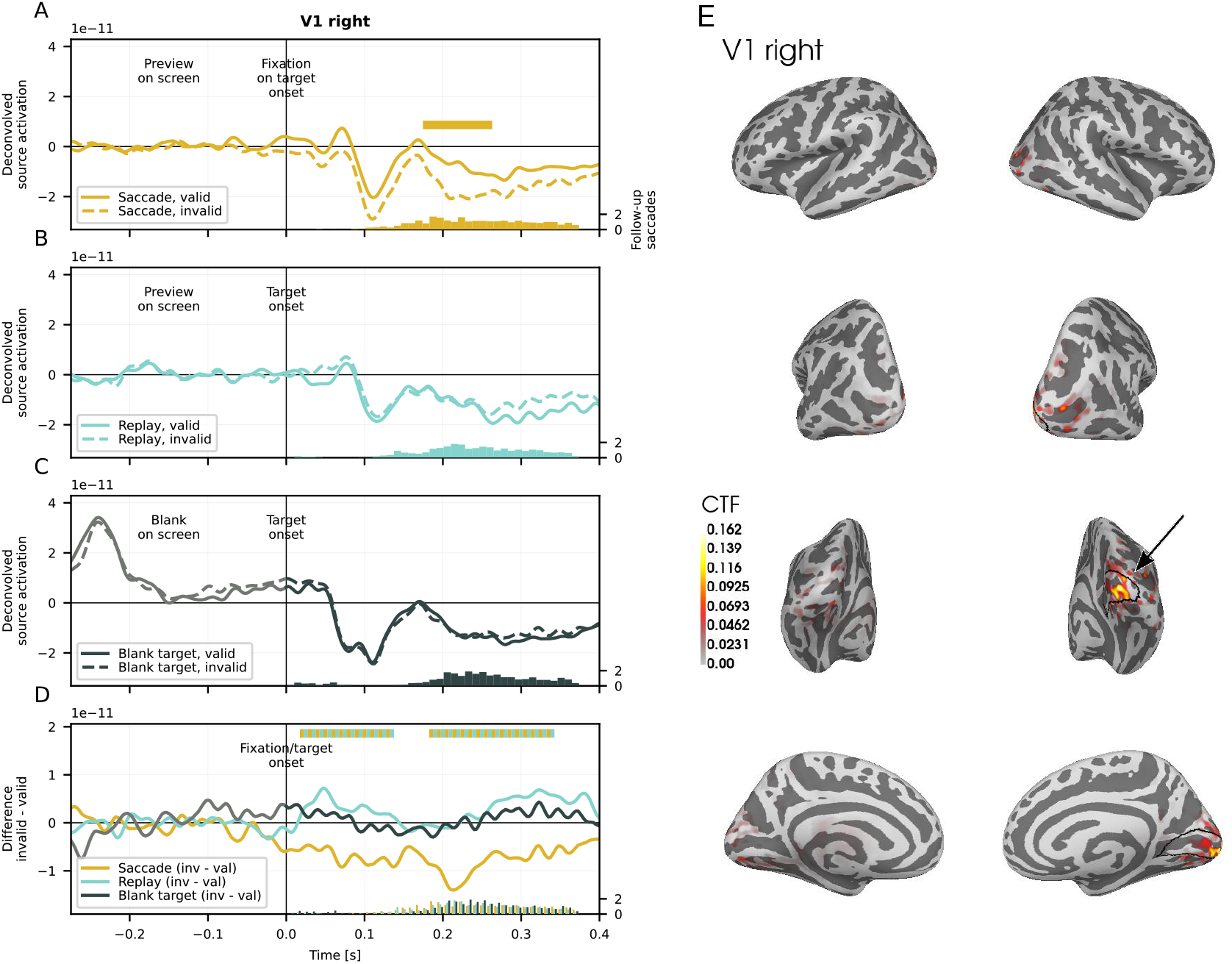
Grand average source-localized and temporally deconvolved MEG signal in the primary visual (V1, left panels) cortex of the right hemisphere and the cross-talk function for this ROI (right panels). (A) Fixation-locked responses in the valid and invalid preview conditions for the active saccade blocks, (B) target-locked responses for the passive replay, and (C) the target-locked responses for the blocks with the blank screen between preview and target stimuli. (D) compares the preview effect contrast invalid minus valid across the three types of viewing blocks. Horizontal bars indicate significant cluster effects: There was a preview effect in the saccade condition which was significantly different from the preview effect in the replay condition (interaction effect clusters in panel D). The contrast comparing the saccade preview effect to the preview effect in blocks with a blank screen between preview and target stimuli did not show any significant clusters. The histograms along the x-axis count the number of follow-up saccades per 10 ms time bins, averaged across participants. (E) Inflated fsaverage anatomy illustrating the cross-talk function (CTF) leakage from all brain sources to the right V1 ROI (black outline), demonstrating that right V1 activity was mostly driven by sources within that ROI.

All other ROIs, apart from left V2, did not show any significant interaction of the preview effect during saccade compared to replay blocks (Figures S1 and S2, all cluster p-values > .05). Left V2 showed a very early viewing (saccade, replay) x preview interaction effect with an opposite sign compared to the early significant cluster in right V1 (Figure 4). These opposite signs might indicate a common neural source but the effect is too early to be related only to the target stimulus because it takes some tens of milliseconds for neural activity from the retina to arrive at V1 ^62,63^. Importantly, however, these early clusters were not accompanied by temporally corresponding preview effects within the saccade or replay viewing conditions and they were absent in an event-locked analysis without deconvolution (Figures S3 and S4). These early interaction clusters in right V1 and left V2 do, thus, not allow any strong conclusions about a modulation of trans-saccadic viewing history effects by active vision. Instead our main finding is that the preview effect was modulated by active vision around 200 ms after target onset in right V1 (Figure 3).

**Figure 4.**
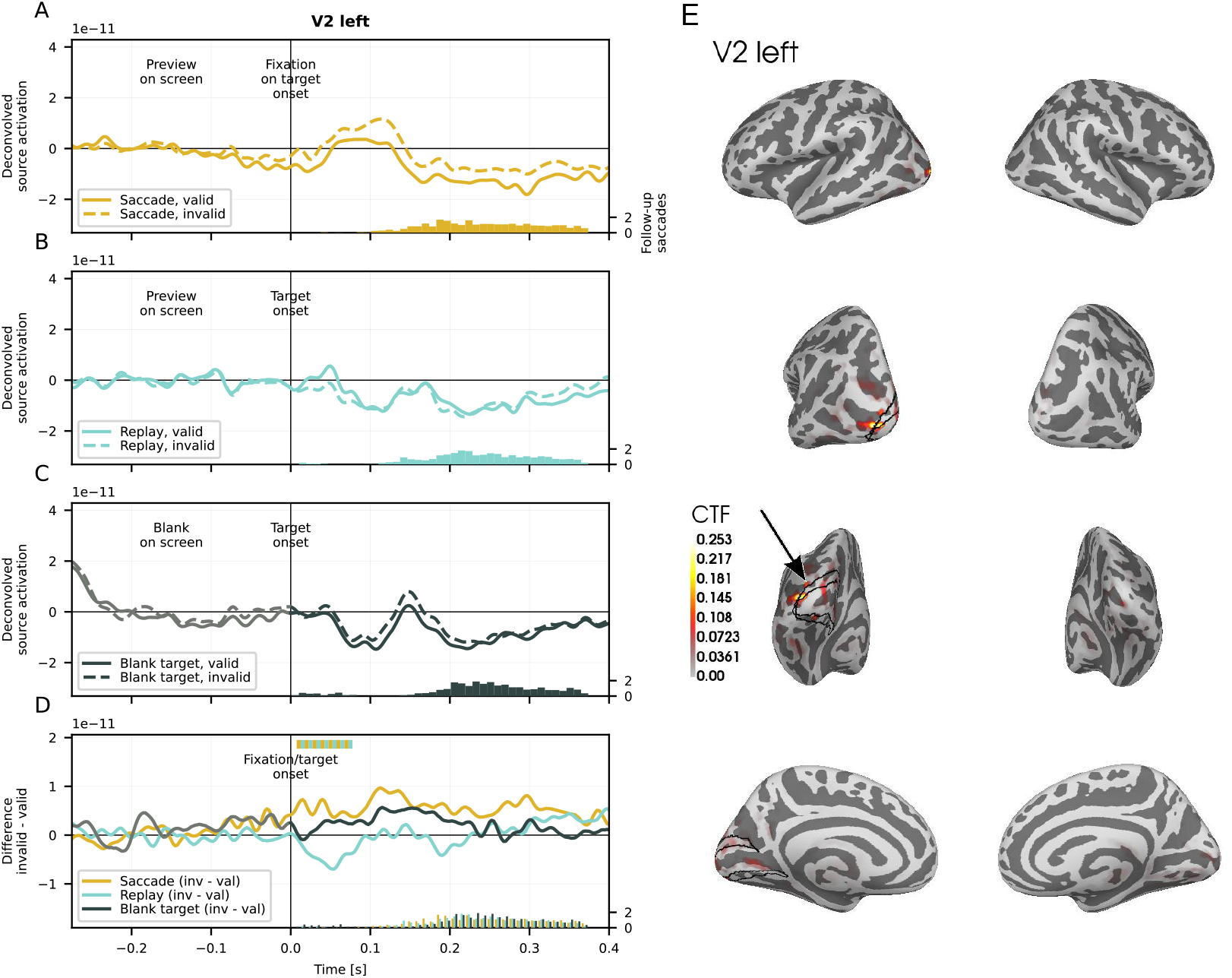
Grand average source-localized and temporally deconvolved MEG signal in left V2. (A) shows fixation-locked responses in the valid and invalid preview conditions for the active saccade blocks, (B) the target-locked response for the passive replay, and (C) the target-locked response for the blocks with the blank screen between preview and target stimuli. (D) compares the preview effect (contrast invalid minus valid) across the three types of viewing blocks. Horizontal bars indicate significant cluster effects: There was one very early significant cluster indicating that the preview effect was different between saccade and replay blocks. In contrast, the preview effect in the saccade compared to the blank blocks was not significantly different. (E) Inflated fsaverage anatomy illustrating the cross-talk function (CTF) leakage from all brain sources to the left V2 ROI (black outline), demonstrating that left V2 activity was mostly driven by sources within that ROI.

The right-lateralized modulation of the preview effect by active vision around 200 ms after target onset is not unexpected considering that the preview stimulus before saccade onset was always in the left visual field and this stimulus was different between valid and invalid preview conditions. In contrast, the right visual field contained a uniform background during the preview phase of a trial (Figure 1A) which was the same for both valid and invalid preview conditions. Thus, in contrast to the right hemisphere, there was no difference in stimulation between valid and invalid preview conditions for the left hemisphere, which can explain why the preview effect was more prevalent in the right hemisphere.

To assess the influence of the deconvolution approach on our results, we repeated the analysis in an event-locked way without deconvolving overlapping evoked potentials triggered by subsequent events and the pattern of results was very similar. As in the deconvolution analysis, the preview effect was more pronounced in saccade than in replay blocks in right V1 around 200 ms after target onset (Figure S3), however the significant interaction cluster was limited to the time period of the significant preview effect within the saccade blocks. The only other ROI where the preview x viewing (saccade, replay) interaction was significant in this event-locked analysis was left SMG with a cluster around 300 to 350 ms, however, without a corresponding preview effect within the saccade or replay conditions (Figure S5), which limits any conclusions from this isolated finding. Moreover, the effect in left SMG was not significant anymore when the two participants with a template brain anatomy were excluded from the analysis whereas our main finding in right V1 was still the same. In sum, the similar influence of active vision on trans-saccadic viewing history effects in the deconvolved compared with the event-locked analysis provides additional support that the viewing history effects in the source-localized MEG signal are not driven by systematic variations in follow-up saccades and their overlapping visuomotor evoked potentials.

### Adaptation alone cannot explain the viewing history effects in visual cortices

Comparing the viewing history effect, that is the preview effect, in the blank blocks to the saccade blocks showed that the influence of active vision on the preview effect in right V1 could not be attributed exclusively to spatiotopic adaptation. Spatiotopic adaptation should have increased the preview effect in the blank blocks compared to the saccade blocks ^48^, but we did not find any evidence for that in right V1 (Figure 3D, all preview (valid, invalid) x viewing block (saccade, blank) interaction contrast cluster p > .05). Moreover, the preview effects in the blank blocks in right V1 were even numerically smaller and not significant compared to the corresponding preview effects in the saccade blocks (Figure 3C). This pattern was exactly the same in an event-locked analysis without deconvolution.

In contrast to right V1, right and left V34 showed a significantly different preview effect in saccade compared to blank blocks with an interaction cluster after 300 ms (Figure S1 and S2), albeit without temporally corresponding significant preview effects within the saccade or blank blocks. This marginal finding could be in line with the spatiotopic adaptation effects found with MRI in overlapping ventral occipital areas that likely overlap with our area V34 ^47,48^. However, if spatiotopic adaptation was the only mechanism for the viewing history effects observed with our preview design in right V1, we would have expected, first, the corresponding interaction effect with the blank blocks also in right V1 and not in V34, and second, this interaction effect should have occurred around the time of the preview effect within the saccade blocks (Figure 3A). Moreover, the interactions with the blank block in right and left V34 were not significant anymore when the two participants with a template brain were excluded from the analysis but our main finding in right V1 was the same.

The event-locked counterpart of the deconvolution analysis revealed additional modulations of the preview effect by the blank blocks in right and left VO and left SMG, however, without temporally corresponding preview effects within saccade or blank blocks (Figures S5 and S6). Apart from this inconsistency, the leakage analysis of right and left VO suggested that these clusters might not be very much confined to VO but rather have a distributed ventral-medial origin (Figure S7). This spatial imprecision unfortunately complicates any firm conclusions from these ROIs and renders right V1 the more reliable result.

Besides spatiotopic adaptation, retinotopic adaptation can also not completely explain the preview effect. Most retinotopic neurons in the visual system will have receptive fields that are too small to be stimulated by both the extrafoveal pre-saccadic preview and the foveal post-saccadic target stimulus, but some neurons might have receptive fields that are large enough to cover both. Receptive fields in the inferotemporal cortex (IT) of macaque monkeys show large variation from a few degrees up to 30-40° ^64–66^. In humans, evidence from population receptive field mapping using fMRI suggests receptive field sizes of 2-12° ^67; in particular Figure 6^ or 4-8° ^68^ in lateral occipital and ventral visual areas. However, if the adaptation of neurons with large receptive fields that covered both the preview and target locations – retinotopically separated by 8° – explained the preview effect, the preview effect should have been the same in saccade and *replay* conditions because in both conditions the stimuli projected to the same retinotopic receptive fields (Figure 3).

### No modulation by active vision in parietal areas

Parietal brain areas typically contain neurons that show transient shifts in retinotopic receptive field locations around the time of saccades, also called predictive remapping if these shifts occur before saccades ^69^ and we did not find any significant preview x viewing interaction effect in intraparietal areas (IP) or in the SMG (Figures S1 and S2, all cluster p-values > .05). This finding reassures that our experimental contrasts worked as expected because they were designed to isolate the influence of a viewing history effect separately for active and passive vision conditions. Although the preview stimulus was probably remapped around the time of saccade execution, we did not expect a difference in this remapping process between valid and invalid preview stimuli, and without such a difference in remapping between valid and invalid preview stimuli, there was no reason to expect that the valid-invalid preview contrast was modulated by remapping. It is, however, not unlikely that parietal areas, or other areas more recently associated with saccadic remapping like V2 ^70^ or V4 ^71^, contributed to trans-saccadic viewing history effects by relaying the efference copy signal from oculomotor to visual areas.

## Discussion

We found that visual cortices exhibit sequential viewing history effects and these sequential effects are modulated by active vision. To determine sequential effects, we used a preview paradigm which has previously been used to demonstrate the impact of pre-saccadic extrafoveal visual input on post-saccadic foveal processing in behavioral measures ^37,41,42^ and in the EEG ^34–36,38–40^. Participants performed a gaze-contingent perceptual task in which they made saccades to extrafoveally presented objects that could change (invalid preview) or remain the same (valid preview) across the saccade. We found a robust preview effect in early visual cortices under this active vision condition, and this preview effect was significantly reduced in V1 and even abolished in ventral-occipital cortices in passive replay blocks where the extrafoveal object moved to the foveal field of view while the participants maintained stable gaze. Such an effect of active vision could in principle be confounded with differences in visual stimulation between the saccade and the replay blocks. However, our preview effect design had the advantage that any such stimulation differences would have cancelled out in the contrast of invalid versus valid preview trials within each of the saccade and replay blocks.

Our preview effect design might have been key to finding sequential viewing history effects specific to active vision. Previous studies either ignored pre-saccadic extrafoveal previews ^14,15,24,25^, or they were lacking either a passive no-saccade replay condition with imitates the visual input elicited by saccade execution ^27^ or an invalid preview condition ^15–17,26^. Importantly, because we contrasted valid and invalid previews to measure sequential viewing history effects, any residual visual stimulation differences that might have existed between our active saccade and passive replay blocks canceled out. To our knowledge, this is the first study that finds sequential viewing history effects in visual cortices that are specific to active vision while ruling out potential confounds from differences in visual stimulation across active and passive viewing conditions.

The same logic of a preview effect design with active and passive viewing conditions has recently been applied by Zhang and colleagues ^45^ to direct recordings from the macaque superior colliculus with remarkably similar results to ours. In that study, macaques made saccades to gratings that could change their spatial frequency (invalid preview) or remain the same (valid preview) during the saccade. In invalid preview conditions, single neuron and population activity increased after fixation onset only if the saccade had been actively made and not if the monkey had maintained fixation and the grating had moved from the extrafoveal to the foveal location. Importantly, this effect was recorded from receptive fields that did not overlap the extrafoveal but only the foveal target location which means that these neurons responded to stimuli that they had not been exposed to ^45^. Our finding shows that such sequential effects are not limited to subcortical brain structures but are present in visual cortices as well.

Crucially, the difference in activity that we found in the visual cortex for a valid compared to an invalid preview, i.e. the preview effect, cannot be explained by neural adaptation ^e.g. 46,72^, repetition suppression ^73,74^, or visual mismatch responses ^75–77^. If some sort of repeated visual stimulation by successive preview and target stimuli was the only mechanism for a preview effect, the preview effect should have been the same in saccade and replay conditions, because in both conditions the same retinotopic neurons were stimulated by both preview and target stimuli. Exclusively spatiotopic adaptation also seems an unlikely explanation, given that spatiotopic adaptation increases with increasing time between adaptor and test ^48^, and inserting a blank interval between our adaptor (preview) and test (target) stimuli rather reduced the preview effect.

It is, however, still possible that a particular type of adaptation in combination with saccadic remapping explains the modulation of sequential effects by active vision. Traditionally, parietal areas, like the lateral intraparietal area (LIP) in monkeys ^57,58^, contain neurons that remap their receptive fields to match the location of upcoming stimuli in the face of an impending saccade ^69,78–82^. This phenomenon of predictive remapping has more recently also been reported for earlier visual areas like V2, V3A, and V4 ^70,71,83^, but we did not find any significant modulation in these areas, nor in the SMG which is also relevant for trans-saccadic perception in humans ^59^. Instead, we found modulations by active vision exactly in the primary and ventral occipital cortices which do not appear to have remapping neurons ^27,83^. It could, thus, be that these visual cortices receive a saccadic efference copy from typical remapping-related areas further downstream which transfers neural activity elicited by the extrafoveal preview in time with the saccade to neurons with foveal receptive fields that cover the post-saccadic target, leading to a post-saccadic adaptation effect at the retinal target location. Crucially, such remapped adaptation cannot result from neural fatigue ^84^, has to act at the timescale of the preview effect of around 200 ms, and relies on a surprisingly fast neural activation transfer within the duration of saccades. This type of remapped adaptation could be the neural basis for rapid perceptual adaptation effects across saccades, which are sometimes known as transiently spatiotopic or craniotopic adaptation ^47,85–87^.

There are several possible neural mechanisms for remapped adaptation. Neural activity could be transferred across retinotopic neural populations through relays to subcortical structures, through areas further downstream ^88^, or horizontally within the visual hierarchy directly across retinotopic cortices. All three possibilities of how retinotopic neural activity could be transferred across saccades make different predictions in terms of bottom-up and top-down information transfer which implies contrasting predictions regarding laminar activation profiles. Bottom-up connections to pyramidal neurons arrive primarily at supragranular layers (2 and 3) whereas top-down connections arrive at infragranular layers (5 and 6) ^89–91^. Evidence from laminar recordings in monkey V1 suggest that fixation onsets are not followed by the typical bottom-up laminar activation profile ^32^, which suggests that some top-down signals arrive at V1 contingent on saccades. This is one potential mechanism that could generate saccadic-specific viewing history effects.

Apart from alterations in bottom-up and top-down processing, the crucial computational principle by which active vision modulates the preview effect could also be of a temporal nature. Temporal expectations are known to enhance effects of expectations ^e.g. 92^ and one might interpret the preview effect as an effect of expectations with a valid preview entailing an expected target in contrast to an invalid preview entailing an unexpected target. Although we made the target temporally predictable also in the passive replay blocks by including a fixed delay between cue and replay onset, the target might still have been more temporally predictable in the saccade blocks simply because saccade execution directly yields the target onset.

The likely contribution from top-down signals related to the efference copy and the consideration about temporal expectations suggests that the saccade-specific viewing history effects we found in early visual cortices might eventually be conceptualized as resulting from a predictive processing mechanism ^5,33,93,94^.

## Conclusion

In sum, our results demonstrate that active vision leads to sequential viewing history effects across saccades in early visual cortices which cannot be explained by neural adaptation alone. Using coregistration of MEG and eye-tracking and harnessing the preview effect in a gaze-contingent experimental protocol we show a larger impact of pre-saccadic extrafoveal input on post-saccadic processing when eye-movements bring objects into focus compared to when the same objects appear during steady fixation. Our study, therefore, provides an update on the role of active vision in modulating cortical function which might be relevant for more naturalistic experimental designs in which eye movements are unavoidable ^95,96^.

## Materials and Methods

### Participants

Coregistered MEG and eye-tracking data was collected from 39 participants. Written informed consent was obtained from all subjects prior to participating in the study which had been approved by the local ethics committee (CMO region Arnhem/Nijmegen). The data of three participants had to be excluded; for one participant because of problems with the head localization coils, for another because of a magnetic artifact, and for one participant because only a fourth of the whole dataset could have been included for the deconvolution analysis based on the trial exclusion criteria described below.

### Stimuli

Eighty objects were selected from the stimulus set of ^97^, available at https://bradylab.ucsd.edu/stimuli/ObjectsAll.zip, considering that the objects should have horizontal or vertical elements to be useful for the main behavioral task, which was a tilt discrimination task. Additional 10 objects were selected for practice trials. The image background was rendered transparent and, if necessary, the original object orientation was adjusted with the image editor GIMP (https://www.gimp.org) to create an upright view of the object. Images were converted to greyscale with the Python module PIL (https://pypi.org/project/pillow/). Subsequently, mean image luminance across all non-background pixels was reduced to 1/8^th^ of the maximum possible luminance, i.e. pixel value 32 out of 255, and pixel luminance standard deviation was set to 16, with the help of the SHINE toolbox ^98^.

For each object, we created a phase-scrambled version which served as invalid preview. Note, that we did not maintain the background-foreground distinction for the scramble object, because this would have made the object easier to recognize, but created a two-dimensional gaussian window in the alpha channel that blended the phase-scrambled object with the background and made the object’s outline harder to recognize. Finally, the luminance of the scrambled images was adjusted to the same value as the luminance of the intact images. All image processing code is available with the experiment code (see section Data and code availability).

The objects had a size of 4° visual angle and were placed with their center at 8° eccentricity to the left of the screen center. Stimulus size and location in visual angels were keep constant across participants by automatically adjusting actual size for each participant individually based on the actually measured screen-eye distance.

A digital light processing (DLP) projector (PROPixx; VPixx Technologies, Saint-Bruno, QC, Canada) rendered the visual stimuli at a refresh rate of 120 Hz on a translucent screen placed 80 cm before the subject. To precisely monitor stimulus onsets, a photodiode was placed in the bottom-right corner of the screen. At that location, a white square was presented with the onset of each stimulus to create a light flash which was recorded via the photodiode into a MISC channel of the MEG system. This procedure revealed the well-known delay of one frame between MEG stimulus onset triggers and the actual change on screen and the time stamps of all stimulus onset triggers events were offline adjusted by the amount of that delay.

### Procedure

The temporal sequence of events within a trial was similar to previous gaze-contingent experiments with the preview effect using face images ^35,44^. In the present study, the experiment consisted of three blocked viewing conditions: a *saccade*, a *replay*, and a *blank* condition (Figure 1). The Participants were informed about which viewing block was coming up at the start of each block. In each of these viewing blocks, a trial started with the presentation of a small fixation dot of 0.1° visual angle presented at screen center and an equally sized placeholder dot located at 8° eccentricity to the left. Stable gaze at screen center for 500 ms triggered the onset of the preview object which had a maximum diameter of 4° and could be either an intact object (*valid* preview) or its phase-scrambled and Gaussian-blurred version (*invalid* preview). The preview object was tilted to the left or right by 2°, which was decided randomly in each trial. Stable gaze was defined in terms of the fixation-update events provided online by the Eyelink eye-tracker. A fixation-update event contains the average gaze position across 50 ms of individual gaze samples. If this position was within 2° visual angle from screen center, it counted as stable gaze. If the participants moved their eyes out of this area, the stable-fixation timer restarted. After 500 ms of stable fixation with the preview object on screen, the saccade cue appeared which consisted in the fixation dot turning slightly darker (Figure 1A).

In the saccade blocks (Figure 1B), participants were instructed to monitor the centrally presented dot and as soon as they noticed the saccade cue they had to look at the extrafoveally presented object. The moment at which the eyes started to move was detected online through a heuristic by calculating the difference in screen pixels between two subsequent gaze samples. If this difference was greater than 8 pixels, it counted as a saccade. This threshold value was determined in pilot testing for this specific lab setting. The detected saccade onset triggered a transient image at the target location which was the scrambled and blurred version of a randomly chosen different object. If the eyes had moved too early, i.e. before the saccade cue, the current trial was appended at the end of the program’s trial queue and the next trial started. The transient image was presented for one frame only and its purpose was to equalize the amount of visual change between valid and invalid conditions. The transient image was followed by the target object which was always an intact object.

The blank blocks were the same as the saccade blocks, expect that instead of the target object a blank screen containing only the placeholder and fixation dots was presented for 400 ms (Figure 1D). Thus, in this condition, the participants saw a blank screen after the saccade. They were instructed to maintain their gaze at the peripheral placeholder location until the target was presented.

The replay blocks were different from the saccade blocks in terms of gaze behavior (Figure 1C). In that condition, participants were instructed to maintain fixation at the screen center throughout the experiment which was controlled by the eye-tracker with the same stable-gaze and saccade detection criteria mentioned above for the saccade blocks. If fixation was too far from screen center or if a saccade was detected, the trial was appended at the end of the trial queue and the next trial started. To match visual input as well as possible to the saccade blocks, the extrafoveal preview object started to move towards the screen center after a fixed time period which was determined for each participant from the saccade latencies that were recorded during the initial practice trials (see below). We call this the *replay* or *replay sequence*. We took the median saccade latency across saccade and blank blocks and subtracted 50 ms to account for the minor speed-up in saccadic latencies that took place throughout the experiment. This adjustment was based on pilot data and previous studies ^35,37^. The preview object moved in equally spaced steps, one per frame, from the extrafoveal preview location to the screen center. The number of frames for this replay sequence was determined based on how many complete frame durations could fit into a participant’s median saccade duration calculated from the practice trial data. The median saccade duration was not adjusted because it does not change much throughout an experiment. The first couple of frames of the replay sequence contained the preview object and only the last frame of the replay sequence contained the transient image to account for the delay between saccade detection and transient presentation in the saccade and blank blocks which also contained only one frame of the transient image. After the replay sequence, the target object was presented at screen center where the participants were still foveating. As in the other viewing blocks, the target object was always intact, that is, not phase scrambled.

Introducing a proper replay sequence in the replay blocks was important in order to minimize visual stimulation differences compared to the saccade blocks because, although we are usually not aware of intra-saccadic visual input, the visual system clearly processes visual input from within a saccadic eye movement ^11^. However, it is in practice impossible to perfectly mimic the visual input arising from a saccadic eye movement. This would require rotating the whole world around a resting eye. To deal with any remaining differences between saccade and replay blocks, our experiment was designed to cancel out any such remaining differences through the preview effect contrast.

In all three viewing blocks, as soon as the participants had foveate the target object, they had to indicate whether it was tilted left (counter-clockwise) or right (clockwise). The tilt of the post-saccadic target object was 2° and always the same for target and pre-saccadic preview object (illustrated in Figure 1B). In all three viewing blocks, the target object disappeared 800 ms after its onset. Manual responses were given with MEG-compatible button-boxes connected to a DataPixx device (VPixx Technologies, Saint-Bruno, QC, Canada).

Each of the 80 objects appeared once in each condition for each participant. With two preview (valid, invalid) and three types of viewing blocks (saccade, replay, blank), this design resulted in 480 trials which were administered in 15 blocks of 32 trials each. Preview condition trials occurred randomly intermixed within each block. The viewing condition was blocked and the order of blocks was pseudorandom in the sense that before a viewing condition could be repeated both of the other two viewing conditions had to have occurred at least equally often. This constraint was set to balance sequence and practice effects that can result from repeated presentation of the same type of contextual condition in trans-saccadic perception studies ^13,cf. 37^. Between viewing blocks, participants could take self-paced breaks. If necessary, the experimenter could also halt the experiment during the task, had the participant moved or had the eye-tracker to be recalibrated.

At the start of an experimental session, the participant received a step-by-step explanation of the sequence of event within a trial and they were walked through one trial per viewing block as example. Afterwards they conducted practice trials in each viewing block with the replay condition being the last one in order to obtain the gaze parameters for the replay sequence from the preceding saccade and blank blocks. During these practice trials, participants received feedback about the correctness of their manual responses in the object tilt discrimination task. If the response was incorrect, the placeholder dot turned red for 300 ms. This feedback was omitted during the experiment proper.

The experiment concluded with one minute of a separate gaze task which had the purpose to obtain MEG data free from task-relevant visual input. This data was added to the training data for the ICA which was conducted to attenuate artifactual eye-movement components primarily related to eyeball motion (see section MEG and eye-tracking data processing and deconvolution analysis, Preprocessing). During this gaze task participants looked back and forth between the fixation dot at screen center and the placeholder dot to the left of fixation. Their gaze was cued by the placeholder and fixation dots alternatingly turning slightly darker in the same way as the saccade cue in the proper experiment.

### MEG and eye-tracking data recording and synchronization

MEG data was collected with a 275-channel CTF system with axial gradiometers in an electromagnetically shielded room at a sampling rate of 1200 Hz. Throughout the recording, the participants’ head position was monitored online with the help of three positioning coils which had been placed at cardinal landmarks at the nasion, left ear, and right ear according to the standard operating procedure in the lab. If the participant had moved too far away from their starting position, they were verbally guided back into their original position in the break between experimental blocks ^99^. In addition to MEG data, horizontal and vertical EOG was collected for the first couple of participants in order to rule out gaze data synchronization issues between MEG and eye-tracking computers. After the experimental session, a set of 250 to 300 head shape points was recorded with a motion tracking system. These head shape points served the offline coregistration of the MEG and the MRI coordinate system.

An Eyelink 1000 plus eye-tracker (SR research, Ontario, Canada) recorded eye gaze concurrently to the MEG at a sampling rate of 1000 Hz in the default eye-tracker data file format (SR research’s EDF). In addition, the gaze data from the eye-tracker was streamed online via an analog output into dedicated MISC channels of the MEG system. Eye gaze events (fixation start/end, saccade start/end, blink start/end) were provided by the SR research gaze parsing algorithm with a saccade velocity threshold of 35 deg/s and a saccade acceleration threshold of 9500 deg/s^2^.

To synchronize MEG and eye-tracking data, trigger events were sent from Psychopy via the python module serial (https://pyserial.readthedocs.io/en/latest/pyserial.html) and a splitter cable connection to the MEG computer and the eye-tracker at the same time. These events were used to offline merge the MEG data with the eye-movement events that had been parsed by the SR research algorithm. The eye-movement events were then used for temporal deconvolution to obtain eye-fixation-locked and stimulus-locked MEG responses.

### Anatomical MRI processing

Anatomical T1-weighted MRI data for each participant was either already available at the institute or had been obtained after the MEG experiment with the exception of two participants. For these two participants, we used freesurfer’s *fsaverage* template brain, skull, and skin meshes warped to the individual participants’ head shape points. Apart from using the anatomical templates all other processing steps were the same as for the other participants. The T1 images were automatically segmented and cortical as well as skull and skin surfaces were reconstructed with freesurfer (version 7.4.1, command) and fastsurfer (https://github.com/Deep-MI/FastSurfer). To ensure that the template brains did not lead to any spurious result, we recomputed the whole analysis without these two participants. The results were the same in this control analysis, unless mentioned in the main text.

According to the mne python standard processing pipeline, the outer skin surface model was used to coregister the anatomical MRI data with the MEG sensor locations by matching the locations of the three fiducials (nasion, left, right ear) and the head shape points to the outer skin surface with an Iterative Closest Point (ICP) fitting algorithm. Manual quality inspection ensured that the coregistration process did not suffer from head shape point outliers or other geometrical abnormalities.

### MEG and eye-tracking data processing and deconvolution analysis

#### Preprocessing

Both the MEG and eye-tracking data was processed with mne python (version 1.6) ^100^. The raw MEG signal had been visually inspected for channels with aberrant signals during data collection and any such bad channels were removed and interpolated with spherical spline interpolation during offline data analysis. A third-order gradient compensation was applied to the MEG data before it was filtered with a 0.1 Hz high-pass and a 40 Hz low-pass Hann-windowed finite impulse response filter using mne python’s default filter option arguments.

In order to remove heartbeat as well as corneoretinal and myogenic eye and eyeblink artefacts in the MEG signal we ran an independent component analysis (ICA) in a separate pipeline which featured an additional 1 Hz high-pass filter to provide the required largely stationary signal for the ICA. An ICA infomax solution was obtained from all successfully completed trials’ data, that is from all trials which had not been aborted because of problems with the trial procedure, and from the separate eye movement task at the end of the experiment. Saccade-related components were automatically labelled by applying the method introduced by ^101^. In short, if the variance ratio of a component’s activation during saccades compared to fixations was larger than 1.1, the component was flagged as saccade related. For most of the participants, visual inspection confirmed the selected components and additionally identified heartbeat components. Three to six components were rejected per participant with the exception of eight components for one participant. The obtained ICA solution weights were then applied to the MEG data in the original pipeline.

In order to integrate the MEG and eye-tracking data in a temporally precise way, the eye-movement events from the Eyelink eye-tracking data file (EDF) were added to the MEG data events array considering the shared experiment triggers which were present in both eye-tracking and MEG data streams ^cf. 102^. Following this procedure, we obtained a maximum coregistration error of 2 ms for all common trigger events. After the coregistration with eye-tracking data events, the MEG data was downsampled to 400 Hz.

We ensured that only data from trials where the participants had properly followed the gaze procedure and where the stimulus onsets had happened in time was selected for further processing. In addition, the response time in the speeded object tilt discrimination task had to be within three median absolute deviations per participant and response option (correct, incorrect). In the saccade and blank conditions, the onset of the participant’s fixation on the target/blank had to be within 50 ms from target/blank onset and not earlier. In the replay condition, any occasional saccades during the constant fixation period had to be smaller than 3°. There was no eye blink from the stable fixation at trial start up until 500 ms after target onset, including the blank period in the blank blocks.

#### Source space transformation

The continuous MEG data was transformed into individual participants source activations with minimum norm estimation (MNE). We used surface source spaces based on the freesurfer white matter segmentation with dipole locations obtained from a recursively subdivided icosahedron downsampled to 5124 locations and a three-layer boundary element model based on the shells between brain, inner skull, outer skull, and outer skin surfaces with default conductivity values of 0.3, 0.006, and 0.3, respectively. Dipole orientations of the source model were restricted to the direction perpendicular to the cortical surface. The noise covariance matrix for the inverse operator was estimated from the baseline period within a trial. For the saccade condition, the baseline period was before the onset of the fixation on the target object. For the replay and blank conditions, the baseline period was before the target onset. The baseline period duration was set to the interval of -275 to -75 ms in order to omit the saccadic spike that was present in the saccade condition. We ensured that all noise covariance matrices were based on the same number of trials per condition in order to avoid that any condition difference in the source localization could have resulted from imbalances in the number of trials.

#### Regions of interest (ROI)

After source space projection, the signal was summarized across the vertices of each anatomical region of interest (ROI). Individual participant’s regions of interest had been obtained according to the multimodal parcellation of the Human Connectome Project (HCP) ^60^. The HCP parcellation, which is available for the freesurfer subject *fsaverage*, was transformed to each participant’s individual anatomical space using the freesufer mri_label2label function. For each individual ROI, the summary source space signal across vertices was computed by finding the dominant orientation across vertices and flipping the sign of vertex activations to match the dominant orientation (see mne python’s *mean_flip* label summary function).

Based on our hypothesis that active vision would modulate visual cortices, we selected a set of ROIs for statistical analyses from the Glasser atlas (Figure 2). The selected ROIs were: The visual areas *V1* and *V2*. Area V3 merged with V4 to obtain a region called *V34* which probably covers parts of the ventral occipital areas relevant for spatiotopic adaptation determined by ^48^. To even better capture activity in ventral-occipital areas, we created an area *VO* from all the remaining ventral-occipital ROIs of the Glasser atlas: V8, VMV3, and the ventral visual complex (VVC). This region VO might be a candidate for neurons with large receptive fields ^65^. The smaller lateral-occipital areas LO1, LO2, LO3 and V4t – V4t is also known as LO2 – were merged to *LO*, because lateral-occipital areas are implicated in object processing and sensory prediction effects ^e.g. 56^ and the current study feature object stimuli. To capture potential effects of saccadic remapping, we added two parietal areas: the intraparietal area *IP* covering the ventral (VIP), medial (MIP) and both ventral and dorsal parts of the lateral intraparietal area (vLIP, dLIP), the intraparietal areas IP1, IP0, and the intraparietal sulcus (IPS1), and the supramarginal gyrus *SMG* consisting of all subparts of Glasser’s parietal area F (PF, PFcm, PFm, PFop, PFt), because that gyrus has been identified as a key region for trans-saccadic updating of object features ^59^ (Figure 2). For further details about the location and differentiation of these areas the reader is referred to the Supplementary Neuroanatomical Results of ^60^.

#### Leakage analysis

To assess the spatial specificity of our ROIs, we conducted a leakage analysis ^61^. For each participant we created the resolution matrix which results from applying the inverse operator to the forward model. We had used minimum norm estimation (MNE) to create the inverse operator which results in a symmetric resolution matrix where the point-spread functions (PSF), how much each source leaks to other sources, and the cross-talk functions (CTF), how much other sources leak to a given source, are equal. Typically, each ROI encompasses a set of source locations and to obtain a one PSF/CTF measure for each ROI, the PSF/CTF was summarized with the help of a principle component analysis (PCA). For each ROI, the first component activation was morphed to the *fsaverage* brain anatomy and averaged across participants. This average leakage was then illustrated separately for each ROI on an inflated *fsaverage* anatomy (Figures 3, 4, and S7).

#### Temporal deconvolution

The source activations summarized per ROI were eventually deconvolved with experiment and eye-movement events using the Unfold.jl toolbox ^52^ (version 0.6.4 and 0.8.11) implemented in the Julia programming language (https://julialang.org, version 1.10.0) and access from within python through the juliacall package (version 0.9.15). Temporal deconvolution requires continuous data at the level of single trials. Sections of the continuous data from in between trials or from trials that were not eligible (see preprocessing above) were set to missing data to exclude them from deconvolution.

We used the maximum number of events for deconvolution given our experimental design, considering that only events which vary in their relative timing across trials can be deconvolved. In line with this constraint, we defined as first event of main interest the onset of the preview image in the saccade and blank blocks. For the saccade block, the second event of main interest was the onset of the first fixation on the target. We did not model the preceding saccade onset separately because the time interval from saccade onset to fixation onset was almost constant. In the blank blocks, the next event of interest was the onset of the blank screen. However, the blank screen and the subsequent target onset had a fixed timing which meant that only one of the events could have been selected for the deconvolution. We selected the target onset because our main hypothesis was about the response to the target stimulus. However, the event basis functions were defined for a time range that was long enough to encompass the onset of the preceding blank screen to additionally obtain insights into potential neural omission responses triggered by that screen onset. Similarly, in the replay blocks the onset of the first event of interest, i.e. the preview object, and the onset of the second event of interest, i.e. the onset of the target, occurred at a fixed timing. This fixed timing served the purpose to render the target temporally predictable because a temporally predictable target was expected to enhance any potential preview effects in this passive replay condition which would make our comparison to the active saccade condition more conservative. For the deconvolution model, this important experimental feature meant that we could only include either the preview onset or the target onset. We opted again for the target onset because the response to the target was of main interest and the deconvolution model was extended in time far enough to encompass the preview onset in order to also compare the neural responses to the preview onsets between all conditions.

Besides the preview onset and the fixation/target onset, we defined four other types of events in the deconvolution model in order to correct for the overlap of their corresponding neural signals: Follow-up saccades after target onsets, additional saccades during the replay sequence which had to be smaller than 3°, additional saccades during presentation of the blank, and the event of the manual response. Participants showed substantial individual variation in the numbers of these three types of additional saccades which happened very likely unintentionally despite the strict gaze-contingent experimental procedure.

The model formulas for each event consisted in the same two predictors preview condition (valid, invalid) and viewing condition (saccade, replay, blank), with the exception of the occasional saccades during the replay sequence and during presentation of the blank screen because these two types of events only occurred within their respective viewing condition and therefore the viewing condition factor was omitted from their model formula. All saccade event formulas and the fixation onset events included saccade amplitude as tenth order spline predictor to account for the non-linear effect of saccade amplitude on the post-saccadic evoked potential ^52^. To make the replay condition comparable to the saccade condition despite the lack of an actual saccade, we included the simulated saccade amplitude of 8 dva as continuous predictor for the target onset event in that condition.

All events were modelled with finite impulse response basis functions but the temporal extent of these basis functions, also known as estimation time window length, varied across event types in line with the time point at which an event could occur within a trial, additionally considering that we wanted to capture all possible overlapping responses. The preview onset time window was -275 ms to 2,000 ms after, the fixation/target onset time window was -1,495 ms to 800 ms, for follow-up saccades, saccades during the replay sequence and during blank presentation it was -275 ms to 800 ms, and for the manual response it was -875 ms to 800 ms. The time windows were based on the maximum time intervals between time-varying events across all participants.

The deconvolution analysis was conducted separately for each participant.^1^ From the resulting regression coefficients, we calculated the predicted means (using the Julia package *Effects*) per condition, region of interest, and time point. Statistical significance was assessed at the group level by applying cluster-based permutation statistics in the time domain using a significance threshold of p = .05 ^103^ for the two planned interaction contrasts preview (valid, invalid) x viewing (saccade, replay) and preview (valid, invalid) x viewing (saccade, blank).

To control for overfitting, we examined the models’ R2. For deconvolution models, R2 might not be a very informative measure. We, thus, applied a cross-validation approach in which we compared each model’s R2 based on the data to the average R2 obtained across 100 iterations of randomly reshuffling the time stamps of the deconvolution events. To test whether the R2 for an ROI was larger than its cross-validated counterpart, we conducted within-participants t-tests. For one of the participants, the random iterations yielded nine negative R2 samples which are a sign of poor model fit for the corresponding random datasets. However, even if these nine samples or the whole participant’s data were excluded, the model fit and the final results remained the same.

#### Event-locked analysis

Since the deconvolution approach added a lot of complexity, we repeated the analysis in a classic event-locked way. Instead of applying deconvolution in the source space, we epoched the source space data around the fixation onset on the saccade block and around target onsets in the replay and blank blocks with a baseline interval of -275 to - 75 ms excluding saccadic spikes present in the saccade blocks before the fixation onset and calculated averages per participant and condition. Based on these source-level activations, statistical significance was assessed in the same way as in the devonvolution pipeline.

### Preregistration

This study was preregistered (https://osf.io/uxtqn). We eventually deviated from the preregistration in the sense that we did not go into the source space for individual vertices but selected regions of interest. This step was necessary for computational reasons related to temporal deconvolution. Initially, we did not consider deconvolution for the preregistration because, based on previous research ^35,44^, we had assumed to get largely the same types of eye movements in both valid and invalid viewing conditions. However, the invalid preview condition with a heavily degraded object eventually led to slightly different saccade characteristics compared to the valid preview condition. To deal with these different characteristics we employed deconvolution which removes the overlapping effect of neural responses to subsequent eye movement events. With deconvolution, projecting sensor data into source space becomes less straightforward, because to go into source space with minimum norm estimation we need a noise-covariance matrix which has to be estimated from single-trial data. Per definition we do not have any single-trial data anymore after deconvolution, only the effects aggregated across trials per conditions per participant. Deconvolution is a type of temporally extended regression which means that the result are regression coefficients which summarize the effect across single-trial data. To get source space results with deconvolution, we had to run the deconvolution analysis already in the source space. There are two options of running unfold on source-projected data: Run it for each vertex or run it for ROIs with signals summarized across vertices within one ROI. We chose the second option, because running deconvolution on each vertex would have been computationally infeasible.

## Data and code availability

All raw data and the code to generate the results from the raw data can be obtained from the Radboud Data Repository under https://doi.org/10.34973/sqkg-xm80.

## Acknowledgements

We thank Robert Oostenveld, Jan-Mathijs Schoffelen, Britta Westner, and Eelke Spaak for insightful comments and suggestions regarding MEG data analysis and for discussion of the results, and Benedikt Ehinger for advice in using Unfold.jl.

This research was supported by a Marie Skłodowska-Curie Individual Fellowship (No. 846392) and by the Italian Ministry of University and Research PNNR NextGenerationEU program (No. 0000027).

## Author contributions

C.H-H.: Conceptualization; Methodology; Investigation (participant recruitment and data collection); Software (task programming and eye-tracking integration; analysis pipelines); Data curation (MEG preprocessing, eye-tracking synchronization and quality control); Formal analysis; Project administration; Visualization; Writing – original draft; Funding acquisition.

F.d.L.: Conceptualization; Supervision; Project administration; Writing – review & editing.

All authors approved the final version of the manuscript and agree to be accountable for the integrity and accuracy of the work.

## Competing interests

The authors declare no competing interests.

## Footnotes

1 We would have preferred to model the data of all participants in one mixed model with random factors for participants and items. This procedure was, however, computationally infeasible, because for only a single data channel the calculations exceeded the maximum resources available on our compute cluster.

## Notes

https://doi.org/10.34973/sqkg-xm80

### Competing Interest Statement

The authors have declared no competing interest.

### Summary of Updates

Added classic event-locked analysis. Added deconvolution model fit assessment. Added analysis of MEG source leakage and updated results to reflect the corresponding reliability of regions of interest. Added parietal ROIs and corresponding changes in background, results, and discussion sections. Corrected a mistake in ICA component rejection for two participant, which minimally affected the source activity waveforms but did not changed the results. Updated introduction and discussion with respect to clarity and significance of the findings.

## References

1. Findlay, J. M. & Gilchrist, I. D. Active Vision. (Oxford University Press, Oxford, UK, 2003). doi:10.1093/acprof:oso/9780198524793.001.0001.

2. Herwig, A. Linking perception and action by structure or process? Toward an integrative perspective. Neurosci. Biobehav. Rev. 52, 105–116 (2015).

3. Rolfs, M. & Schweitzer, R. Coupling perception to action through incidental sensory consequences of motor behaviour. Nat. Rev. Psychol. 1, 112–123 (2022).

4. Cont, C. & Zimmermann, E. The Motor Representation of Sensory Experience. Curr. Biol. 31, 1029–1036.e2 (2021).

5. Keller, G. B. & Mrsic-Flogel, T. D. Predictive Processing: A Canonical Cortical Computation. Neuron 100, 424–435 (2018).

6. Cavanaugh, J., Berman, R. A., Joiner, W. M. & Wurtz, R. H. Saccadic corollary discharge underlies stable visual perception. J. Neurosci. 36, 31–42 (2016).

7. Melcher, D. Visual stability. Philos. Trans. R. Soc. B Biol. Sci. 366, 468–475 (2011).

8. von Holst, E. & Mittelstaedt, H. Das Reafferenzprinzip [The reafference principle]. Naturwissenschaften 37, 464–476 (1950).

9. Helmholtz, H. von. Handbuch Der Physiologischen Optik [Handbook of Physiological Optics]. (Voss, Leipzig, 1867).

10. Mostofi, N. et al. Spatiotemporal Content of Saccade Transients. Curr. Biol. 30, 3999–4008.e2 (2020).

11. Schweitzer, R. & Rolfs, M. Intrasaccadic motion streaks jump-start gaze correction. Sci. Adv. 7, eabf2218 (2021).

12. Grzeczkowski, L., Shi, Z., Rolfs, M. & Deubel, H. Perceptual learning across saccades: Feature but not location specific. Proc. Natl. Acad. Sci. 120, e2303763120 (2023).

13. Scholes, C., McGraw, P. V. & Roach, N. W. Learning to silence saccadic suppression. Proc. Natl. Acad. Sci. U. S. A. 118, (2021).

14. DiCarlo, J. J. & Maunsell, J. H. R. Form representation in monkey inferotemporal cortex is virtually unaltered by free viewing. Nat. Neurosci. 3, 814–821 (2000).

15. Gawne, T. J. & Martin, J. M. Responses of Primate Visual Cortical Neurons to Stimuli Presented by Flash, Saccade, Blink, and External Darkening. J. Neurophysiol. 88, 2178–2186 (2002).

16. Richmond, B. J., Hertz, J. A. & Gawne, T. J. The relation between V1 neuronal responses and eye movement-like stimulus presentations. Neurocomputing 26–27, 247–254 (1999).

17. Ruiz, O. & Paradiso, M. A. Macaque V1 representations in natural and reduced visual contexts: Spatial and temporal properties and influence of saccadic eye movements. J. Neurophysiol. 108, 324–333 (2012).

18. MacEvoy, S. P., Hanks, T. D. & Paradiso, M. A. Macaque V1 Activity During Natural Vision: Effects of Natural Scenes and Saccades. J. Neurophysiol. 99, 460–472 (2008).

19. McFarland, J. M., Bondy, A. G., Saunders, R. C., Cumming, B. G. & Butts, D. A. Saccadic modulation of stimulus processing in primary visual cortex. Nat. Commun. 6, 8110 (2015).

20. Binda, P. & Morrone, M. C. Vision during saccadic eye movements. Annu. Rev. Vis. Sci. 4, 193–213 (2018).

21. Boi, M., Poletti, M., Victor, J. D. & Rucci, M. Consequences of the Oculomotor Cycle for the Dynamics of Perception. Curr. Biol. 27, 1268–1277 (2017).

22. Nicolas, G. et al. Neural correlates of intra-saccadic motion perception. J. Vis. 21, 19 (2021).

23. Schweitzer, R., Seel, T., Raisch, J. & Rolfs, M. Early visual signatures and benefits of intra-saccadic motion streaks. PLOS Comput. Biol. 21, e1013544 (2025).

24. Gallant, J. L., Connor, C. E. & Van Essen, D. C. Neural activity in areas V1, V2 and V4 during free viewing of natural scenes compared to controlled viewing: NeuroReport 9, 2153–2158 (1998).

25. Kagan, I., Gur, M. & Snodderly, D. M. Saccades and drifts differentially modulate neuronal activity in V1: Effects of retinal image motion, position, and extraretinal influences. J. Vis. 8, 19–19 (2008).

26. Vinje, W. E. & Gallant, J. L. Sparse Coding and Decorrelation in Primary Visual Cortex During Natural Vision. Science 287, 1273–1276 (2000).

27. Xiao, W., Sharma, S., Kreiman, G. & Livingstone, M. S. Feature-selective responses in macaque visual cortex follow eye movements during natural vision. Nat. Neurosci. 1–10 (2024) doi:10.1038/s41593-024-01631-5.

28. Leszczynski, M. & Schroeder, C. E. The Role of Neuronal Oscillations in Visual Active Sensing. Front. Integr. Neurosci. 13, (2019).

29. Schroeder, C. E., Wilson, D. A., Radman, T., Scharfman, H. & Lakatos, P. Dynamics of Active Sensing and perceptual selection. Curr. Opin. Neurobiol. 20, 172–176 (2010).

30. Leszczynski, M. et al. Eye movements organize excitability state, information coding and network connectivity in the human hippocampus. Prog. Neurobiol. 252, 102812 (2025).

31. Barczak, A. et al. Dynamic Modulation of Cortical Excitability during Visual Active Sensing. Cell Rep. 27, 3447–3459.e3 (2019).

32. Rajkai, C. et al. Transient cortical excitation at the onset of visual fixation. Cereb. Cortex 18, 200–209 (2008).

33. Huber-Huber, C., Buonocore, A. & Melcher, D. The extrafoveal preview paradigm as a measure of predictive, active sampling in visual perception. J. Vis. 21, 12 (2021).

34. Buonocore, A., Dimigen, O. & Melcher, D. Post-saccadic face processing is modulated by pre-saccadic preview: Evidence from fixation-related potentials. J. Neurosci. 0861–19 (2020) doi:10.1523/JNEUROSCI.0861-19.2020.

35. Huber-Huber, C., Buonocore, A., Dimigen, O., Hickey, C. & Melcher, D. The peripheral preview effect with faces: Combined EEG and eye-tracking suggests multiple stages of trans-saccadic predictive and non-predictive processing. NeuroImage 200, 344–362 (2019).

36. de Lissa, P. et al. Peripheral preview abolishes N170 face-sensitivity at fixation: Using fixation-related potentials to investigate dynamic face processing. Vis. Cogn. 27, 740–759 (2019).

37. Huber-Huber, C. & Melcher, D. The behavioural preview effect with faces is susceptible to statistical regularities: Evidence for predictive processing across the saccade. Sci. Rep. 11, 942 (2021).

38. Degno, F. et al. Parafoveal previews and lexical frequency in natural reading: Evidence from eye movements and fixation-related potentials. J. Exp. Psychol. Gen. 148, 453–474 (2019).

39. Dimigen, O., Kliegl, R. & Sommer, W. Trans-saccadic parafoveal preview benefits in fluent reading: A study with fixation-related brain potentials. NeuroImage 62, 381–393 (2012).

40. Hutzler, F. et al. Welcome to the real world: Validating fixation-related brain potentials for ecologically valid settings. Brain Res. 1172, 124–129 (2007).

41. Rayner, K. The perceptual span and peripheral cues in reading. Cognit. Psychol. 7, 65–81 (1975).

42. Schotter, E. R., Angele, B. & Rayner, K. Parafoveal processing in reading. Atten. Percept. Psychophys. 74, 5–35 (2012).

43. Kornrumpf, B., Niefind, F., Sommer, W. & Dimigen, O. Neural correlates of word recognition: A systematic comparison of natural reading and rapid serial visual presentation. J. Cogn. Neurosci. 28, 1374–1391 (2016).

44. Huber-Huber, C. & Melcher, D. Saccade execution increases the preview effect with faces: An EEG and eye-tracking coregistration study. Atten. Percept. Psychophys. 10.3758/s13414-023-02802-5 (2023) doi:10.3758/s13414-023-02802-5.

45. Zhang, T., Bogadhi, A. R. & Hafed, Z. M. Foveal neurons of the monkey superior colliculus signal trans-saccadic prediction errors. PLOS Biol. 23, e3003246 (2025).

46. Benda, J. Neural adaptation. Curr. Biol. 31, R110–R116 (2021).

47. Zimmermann, E., Morrone, M. C., Fink, G. R. & Burr, D. Spatiotopic neural representations develop slowly across saccades. Curr. Biol. 23, R193–R194 (2013).

48. Zimmermann, E., Weidner, R., Abdollahi, R. O. & Fink, G. R. Spatiotopic adaptation in visual areas. J. Neurosci. 36, 9526–9534 (2016).

49. Zimmermann, E., Weidner, R. & Fink, G. R. Spatiotopic updating of visual feature information. J. Vis. 17, 1–9 (2017).

50. Melcher, D. & Deodato, M. Predicting what and when across saccades: Evidence from the extrafoveal preview effect. J. Vis. 26, 3 (2026).

51. Dimigen, O. & Ehinger, B. V. Regression-based analysis of combined EEG and eye-tracking data: Theory and applications. J. Vis. 21, 3 (2021).

52. Ehinger, B. V. & Dimigen, O. Unfold: an integrated toolbox for overlap correction, non-linear modeling, and regression-based EEG analysis. PeerJ 1–33 (2019) doi:10.7717/peerj.7838.

53. Smith, N. J. & Kutas, M. Regression-based estimation of ERP waveforms: I. The rERP framework. Psychophysiology 52, 157–168 (2015).

54. Smith, N. J. & Kutas, M. Regression-based estimation of ERP waveforms: II. Nonlinear effects, overlap correction, and practical considerations. Psychophysiology 52, 169–181 (2015).

55. Evans, C. C. Spontaneous excitation of the visual cortex and association areas — Lambda waves. Electroencephalogr. Clin. Neurophysiol. 5, 69–74 (1953).

56. Richter, D., Ekman, M. & de Lange, F. P. Suppressed sensory response to predictable object stimuli throughout the ventral visual stream. J. Neurosci. 38, 7452–7461 (2018).

57. Merriam, E. P., Genovese, C. R. & Colby, C. L. Spatial updating in human parietal cortex. Neuron 39, 361–373 (2003).

58. Duhamel, J.-R., Colby, C. L. & Goldberg, M. E. The updating of the representation of visual space in parietal cortex by intended eye movements. Science 255, 90–92 (1992).

59. Baltaretu, B. R., Stevens, W. D., Freud, E. & Crawford, J. D. Occipital and parietal cortex participate in a cortical network for transsaccadic discrimination of object shape and orientation. Sci. Rep. 13, 11628 (2023).

60. Glasser, M. F. et al. A multi-modal parcellation of human cerebral cortex. Nature 536, 171–178 (2016).

61. Hauk, O., Stenroos, M. & Treder, M. S. Towards an objective evaluation of EEG/MEG source estimation methods – The linear approach. NeuroImage 255, 119177 (2022).

62. Bullier, J. Integrated model of visual processing. Brain Res. Rev. 36, 96–107 (2001).

63. Inui, K. & Kakigi, R. Temporal Analysis of the Flow From V1 to the Extrastriate Cortex in Humans. J. Neurophysiol. 96, 775–784 (2006).

64. De Beeck, H. O. & Vogels, R. Spatial sensitivity of macaque inferior temporal neurons. J. Comp. Neurol. 426, 505–518 (2000).

65. DiCarlo, J. J., Zoccolan, D. & Rust, N. C. How does the brain solve visual object recognition? Neuron 73, 415–434 (2012).

66. Kobatake, E. & Tanaka, K. Neuronal selectivities to complex object features in the ventral visual pathway of the macaque cerebral cortex. J. Neurophysiol. 71, 856–867 (1994).

67. Amano, K., Wandell, B. A. & Dumoulin, S. O. Visual Field Maps, Population Receptive Field Sizes, and Visual Field Coverage in the Human MT+ Complex. J. Neurophysiol. 102, 2704–2718 (2009).

68. Dumoulin, S. O. & Wandell, B. A. Population receptive field estimates in human visual cortex. NeuroImage 39, 647–660 (2008).

69. Colby, C. L. & Goldberg, M. E. Space and attention in parietal cortex. Annu. Rev. Neurosci. 22, 319–49 (1999).

70. Denagamage, S., Morton, M. P., Hudson, N. V. & Nandy, A. S. Widespread receptive field remapping in early primate visual cortex. Cell Rep. 43, 114557 (2024).

71. Neupane, S., Guitton, D. & Pack, C. C. Two distinct types of remapping in primate cortical area V4. Nat. Commun. 7, 10402 (2016).

72. Weber, A. I., Krishnamurthy, K. & Fairhall, A. L. Coding Principles in Adaptation. Annu. Rev. Vis. Sci. 5, 427–449 (2019).

73. Grill-Spector, K., Henson, R. & Martin, A. Repetition and the brain: Neural models of stimulus-specific effects. Trends Cogn. Sci. 10, 14–23 (2006).

74. Grotheer, M. & Kovács, G. Can predictive coding explain repetition suppression? Cortex 80, 113–124 (2016).

75. Garrido, M. I., Kilner, J. M., Stephan, K. E. & Friston, K. J. The mismatch negativity: A review of underlying mechanisms. Clin. Neurophysiol. 120, 453–463 (2009).

76. Pazo-Alvarez, P., Cadaveira, F. & Amenedo, E. MMN in the visual modality: A review. Biol. Psychol. 63, 199–236 (2003).

77. Stefanics, G., Kremláček, J. & Czigler, I. Visual mismatch negativity: A predictive coding view. Front. Hum. Neurosci. 8, 1–19 (2014).

78. Burr, D. C. & Morrone, M. C. Spatiotopic coding and remapping in humans. Philos. Trans. R. Soc. B Biol. Sci. 366, 504–515 (2011).

79. Neupane, S., Guitton, D. & Pack, C. C. Perisaccadic remapping: What? How? Why? Rev. Neurosci. 31, 505–520 (2020).

80. Sommer, M. A. & Wurtz, R. H. Brain circuits for the internal monitoring of movements. Annu. Rev. Neurosci. 31, 317–338 (2008).

81. Sun, L. D. & Goldberg, M. E. Corollary discharge and oculomotor proprioception: Cortical mechanisms for spatially accurate vision. Annu. Rev. Vis. Sci. 2, 61–84 (2016).

82. Zirnsak, M. & Moore, T. Saccades and shifting receptive fields: Anticipating consequences or selecting targets? Trends Cogn. Sci. 18, 621–628 (2014).

83. Nakamura, K. & Colby, C. L. Updating of the visual representation in monkey striate and extrastriate cortex during saccades. Proc. Natl. Acad. Sci. 99, 4026–4031 (2002).

84. Whitmire, C. J. & Stanley, G. B. Rapid Sensory Adaptation Redux: A Circuit Perspective. Neuron 92, 298–315 (2016).

85. Cicchini, G. M., Binda, P., Burr, D. C. & Morrone, M. C. Transient spatiotopic integration across saccadic eye movements mediates visual stability. J. Neurophysiol. 109, 1117–1125 (2013).

86. Fabius, J. H., Fracasso, A., Nijboer, T. C. W. & Van Der Stigchel, S. Time course of spatiotopic updating across saccades. Proc. Natl. Acad. Sci. 116, 2027–2032 (2019).

87. Fairhall, S. L., Schwarzbach, J., Lingnau, A., Van Koningsbruggen, M. G. & Melcher, D. Spatiotopic updating across saccades revealed by spatially-specific fMRI adaptation. NeuroImage 147, 339–345 (2017).

88. Khayat, P. S., Spekreijse, H. & Roelfsema, P. R. Correlates of transsaccadic integration in the primary visual cortex of the monkey. Proc. Natl. Acad. Sci. 101, 12712–12717 (2004).

89. Bastos, A. M. et al. Canonical microcircuits for predictive coding. Neuron 76, 695–711 (2012).

90. Bonaiuto, J. J. et al. Lamina-specific cortical dynamics in human visual and sensorimotor cortices. eLife 7, 1–32 (2018).

91. Thomas, E. R. et al. Predictions and errors are distinctly represented across V1 layers. Curr. Biol. 34, 2265–2271.e4 (2024).

92. Nobre, A., Correa, A. & Coull, J. The hazards of time. Curr. Opin. Neurobiol. 17, 465–470 (2007).

93. Friston, K., Adams, R. A., Perrinet, L. & Breakspear, M. Perceptions as hypotheses: Saccades as experiments. Front. Psychol. 3, 1–20 (2012).

94. Parr, T. & Friston, K. J. Active inference and the anatomy of oculomotion. Neuropsychologia 111, 334–343 (2018).

95. Stangl, M., Maoz, S. L. & Suthana, N. Mobile cognition: Imaging the human brain in the ‘real world’. Nat. Rev. Neurosci. 24, 347–362 (2023).

96. Yates, J. L. et al. Detailed characterization of neural selectivity in free viewing primates. Nat. Commun. 14, 3656 (2023).

97. Brady, T. F., Konkle, T., Alvarez, G. A. & Oliva, A. Visual long-term memory has a massive storage capacity for object details. Proc. Natl. Acad. Sci. 105, 14325–14329 (2008).

98. Willenbockel, V. et al. Controlling low-level image properties: the SHINE toolbox. Behav. Res. Methods 42, 671–684 (2010).

99. Stolk, A., Todorovic, A., Schoffelen, J.-M. & Oostenveld, R. Online and offline tools for head movement compensation in MEG. NeuroImage 68, 39–48 (2013).

100. Gramfort, A. et al. MEG and EEG data analysis with MNE-Python. Front. Neurosci. 7, 1–13 (2013).

101. Plöchl, M., Ossandón, J. P. & König, P. Combining EEG and eye tracking: Identification, characterization, and correction of eye movement artifacts in electroencephalographic data. Front. Hum. Neurosci. 6, 278 (2012).

102. Dimigen, O., Sommer, W., Hohlfeld, A., Jacobs, A. M. & Kliegl, R. Coregistration of eye movements and EEG in natural reading: Analyses and review. J. Exp. Psychol. Gen. 140, 552–572 (2011).

103. Maris, E. & Oostenveld, R. Nonparametric statistical testing of EEG- and MEG-data. J. Neurosci. Methods 164, 177–190 (2007).

